# Psychopathy and distance estimation

**DOI:** 10.1101/2022.12.16.520720

**Authors:** Georgios Michalareas, Claudia Lehr, Matthias Grabenhorst, Heiko Hecht

## Abstract

Research on psychopathy has so far been largely limited to the investigation of high-level processes, such as emotion perception and regulation. In the present work, we investigate whether psychopathy has an effect on the estimation of fundamental physical parameters, which are computed in the brain during early stages of sensory processing. We employed a simple task in which participants had to estimate their interpersonal distance from a moving avatar and stop it at a given distance. The face expression of the avatars were positive, negative, or neutral. Participants carried out the task online on their home computers. We measured the psychopathy level via a self-report questionnaire. Regardless of the degree of psychopathy, the facial expression of the avatars showed no effect on distance estimation. Our results show that individuals with a high degree of psychopathy underestimate distance of approaching avatars significantly less (let the avatar approach them significantly closer) than did participants with a lesser degree of psychopathy. Moreover, participants who scored high in Self-Centered Impulsivity underestimate the distance to approaching avatars significantly less (let the avatar approach closer) than participants with a low score. Distance estimation is considered an automatic process performed at early stages of visual processing. Therefore, our results imply that psychopathy affects basic early sensory processes, such as feature extraction, in the visual cortex.

## Introduction

The concept of psychopathy has been studied for many decades and is associated with a set of disruptive personality traits and antisocial behavior. Despite these extensive research efforts, it remains still largely unclear which aspects of behavior and which brain processes are affected by psychopathy. One of the most studied aspects in this context are deficits in emotional processing in people with a high level of psychopathy, especially with respect to facial expressions of other people. It has been proposed that deficits in the processing of such cues are an inherent part of psychopathy. Typically, the brain correlates of these deficits are assumed to exist in brain areas related to higher processes, such as recognition of emotions and regulation of response to them. The influence of psychopathy on more fundamental brain processes has so far played a rather subordinate role in the literature. The current work is interested in studying whether there are effects of psychopathy on such fundamental processes. For this reason, we use an interpersonal distance task in which we investigate whether emotional context affects estimation of the most basic parameter, distance itself. Finding behavioural evidence that psychopathy affects automated processes in early visual hierarchy, such as distance estimation, would be a novel finding showing that psychopathy has underlying brain correlates even in sensory cortex.

### Psychopathy

The current research trend assumes that psychopathic personality traits occur at varying degrees of intensity in the general population and are thus not exclusive to subsets of criminal offenders. In 1941, Hervey Cleckley was the first to define the term psychopathy by describing 16 characteristics. According to his concept definition, a highly psychopathic person can be described with characteristics such as superficial charm, irresponsibility, fearlessness, dishonesty, egocentrism and anti-social behavior. Moreover, there is a lack of emotional depth, impaired learning from punishment, foresight and planning, externalized guilt and not being able to value friendliness (Cleckley, 1964).

Later, Robert D. Hare extended the basic assumptions of Cleckley, and developed criteria for assessing psychopathic personality traits (Hare et al., 1990). According to Hare, psychopathy is mainly characterized by impulsive, thrill-seeking behaviors combined with anxiety, dishonesty, egocentricity, manipulation and exploitation of others (Hare, 1991). His research focused mainly on prisoners. For practical application, the so-called Psychopathy Checklist-Revised (PCL-R, Hare et al., 1990) was generated from these criteria. It contains not only the characteristics defined by Cleckley, but also criminal behavior. It is still used in a revised version as a risk assessment tool for offenders. However, the PCL-R is unsuitable for use with non-forensic samples.

Cleckley in contrast did not understand criminal behavior as a central component of psychopathy, as emphasized by Skeem and Cooke (2010). His research included so-called successful psychopaths in the concept of psychopathy (Cleckley, 1976). Although they share the same main characteristics, successful psychopaths can be distinguished from unsuccessful ones by a higher ability to adapt well in society and avoid clashes with the law. From these considerations it is clear that criminal behavior is not necessarily part of the construct and that psychopathic personality traits are not found exclusively in forensic populations.

Many publications on psychopathy report that psychopathy traits are associated with low empathy (e.g. Soderstrom, 2003; Hare, 2006; Blair, 2005). Even more specifically, there is evidence that psychopaths have impaired processing when recognizing emotional facial expressions such as fear (Montagne et al. 2005). People with psychopathic characteristics can be intriguing businesspersons, employees who use their professional position to victimize others and to enrich themselves at their expense. An overrepresentation of psychopathic individuals can be found in certain professions such as politics, law enforcement, firefighting, and risky sports or military combat (Babiak & Hare, 2006; Fowles & Dindo, 2006; Stevens, Deuling, & Armenakis, 2012).

By now, unlike the PCL of Hare, there exist other questionnaires, which are more suitable for assessing a successful psychopathic sample. The Psychopathic Personality Inventory-Revised [PPI-R; (Lilienfeld & Windows, 2005)] excludes items that refer to criminal or antisocial behavior, assuming that criminality is not a central feature of psychopathy. It is a self-report questionnaire designed to measure psychopathic traits in a dimensional manner. It reflects the idea that psychopathy is a set of traits continuously distributed in the general population and not a discrete forensic pathology (Krueger et al., 2005; Skeem et al., 2011). The PPI-R focuses on eight characteristics of psychopathy measured with eight subscales. These subscales, with one exception, load on two higher order factors. Social influence, stress immunity, and fearlessness can be consolidated as “Fearless Dominance”. This factor represents emotional and interpersonal deficits and is also related to positive adjustment, such as charming behavior (e.g., “Even when others are upset with me, I can usually win them over with my charm”). The characteristics machiavellian egocentricity, rebellious nonconformity, blame externalization, and carefree non-planfulness can be summarized as “Self-Centered Impulsivity”. This factor is related to disinhibition and impulsive behavior (e.g., “I like to act first and think later”). Apart from these two main components the PPI-R also includes the characteristic coldheartedness, which is associated with low empathy, lack of compassion and indifference towards other people’s feelings (e.g., “When someone is hurt by something I say or do, that’s their problem”) (Alpers & Eisenbarth, 2008). Although the PPI-R was developed to assess psychopathic personality traits in community samples, the scale correlates with psychopathy assessment instruments that are used for forensic samples, such as the PCL-R (Poythress et al., 2010). The PPI-R has already been used to assess psychopathy levels in student samples (e.g. Welsch, Hecht, von Castell, 2018; Vieira & Marsh, 2014).

### Psychopathy and social distance

Although psychopathy is characterized by severe deficits in social emotional behavior, the exact mechanisms and the level of brain processing involved are still poorly understood. A few studies in the past have focused on the impact of psychopathy on interpersonal spatial behavior (e.g. Welsch et al., 2018; Vieira & Marsh, 2014). This work is typically based on a research field called proxemics, which was pioneered by Hall (1963). It covers the investigation of human use of space and the effect of behavior, communication and social interaction. Hall describes four zones of interpersonal distances that form different types of spaces: Intimate distance for embracing, touching or whispering (up to 46 cm), personal distance for interactions among good friends or family (up to 122 cm), social distance for interactions among acquaintances (up to 370 cm), and public distance used for public speaking.

Interpersonal space can be influenced by various factors like personal preferences or situational context. In addition, interpersonal space is altered by individual factors. Regarding psychopathy, some studies found evidence for altered spatial behavior in individuals with higher levels of psychopathy. For instance, Vieira and Marsh (2014) showed that Coldheartedness, one of the main characteristics of psychopathy, has a significant effect on interpersonal distance preference. Their results demonstrate that those who scored high on the Coldheartedness scale maintained a lower overall interpersonal distance compared with participants who scored low on this scale.

Experiments of Welsch et al. (2018) focused on the impact of psychopathy on preferred interpersonal distance. They sought to find out why people with higher psychopathy levels may violate personal space requirements. Taking into account that nonverbal cues in social situations play an important role on choosing appropriate distances, they decided to examine the effect of different facial expressions as social cues on preferred IPD. Their research was based on prior findings by Ruggiero et al. (2017), who conducted a virtual reality experiment to investigate spatial behavior while varying the facial expressions of virtual persons. Results showed that participants kept a larger distance towards persons with angry facial expressions in contrast to happy or neutral facial expressions. Since reaction to threat is impaired in people with a high psychopathic level (e.g. von Borries et al., 2012), Welsch et al. (2018) hypothesized that they should fail to regulate their interpersonal distance as a function of facial expressions appropriately. They used the PPI-R mentioned above to separate the student sample into two groups: subjects with high vs. low psychopathic traits. Participants were immersed in a virtual environment for a highly controlled set up. The task was to walk towards an avatar until a comfortable distance for conversation had been reached, as if the participant had to ask a stranger for directions. As the main experimental factor, the avatar’s face expression in each trial was either happy or angry, in order to examine spatial behavior in threatening and safe social encounters. Consistent with results of Ruggiero et al. (2017), participants preferred a larger distance to avatars with angry facial expressions in comparison to ones with happy face expression. There was no overall effect of psychopathy, but a significant interaction effect of facial expression and psychopathy was found and investigated in more detail. As hypothesized, for participants with high psychopathy scores, facial expression had a smaller modulating effect as compared to participants with low psychopathy scores. In contrast to the work of Vieira and Marsh (2014), they did not find Coldheartedness to be a significant predictor for a preference of a shorter interpersonal distance. Also the other facets of the PPI-R score, namely Fearless Dominance and Self-Centered Impulsivity, were not associated with shorter preferred distance. But Coldheartedness and Self-Centered Impulsivity equally predicted a diminished effect of facial expressions on interpersonal distance regulation. Welsch et al. (2018) assume that this reflects the tendency of high psychopathic individuals to not integrate properly information from peripheral social cues when engaging in goal-directed behavior. The finding can be explained by the “response modulation hypothesis of psychopathy”. This theory claims that psychopathy is an *attention* disorder that prevents individuals with high levels of psychopathy to shift their attention to peripheral cues as long as they focus on a particular main goal. According to this theory, there is no inherent lack of empathy or fear, but psychopaths just do not focus on emotional cues if they do not relate to their current goal. The authors argue that psychopaths have normal levels of fear and empathy when they are asked to focus on relevant cues (Smith & Lilienfeld, 2015). Subsequently the general understanding of interpersonal spatial relationships, not from a personal but from an observer’s point of view, was also investigated by Welsch et al. (2018). In a second experiment, participants had to adjust the distance between two avatars until a comfortable interpersonal distance for conversation had been reached. Hence, participants were not personally involved in the interpersonal distance. Overall, the results show that high psychopathic participants had the same concept of comfortable distance as low psychopathic participants. Welsch et al. (2018) conclude from their research, that psychopathy is not in general associated with a changed perception of comfortable interpersonal distance. It is rather the case that psychopathic individuals fail to regulate interpersonal distance with respect to facial expression of an approached person.

However, the exact underlying mechanism remains unclear. Notably, in many studies in the field of interpersonal distance preference, the concept of comfort is involved, as participants are either asked to approach an avatar until “a comfortable distance for conversation was reached” (Welsch et al., 2018) or to stop an approaching person at the distance at which they felt “the most comfortable” (Vieira & Marsh, 2014). Processing comfort in the instructions can involve low- and high-level processes. The type of facial expression could affect higher brain processes, involved in emotion regulation, or could affect much lower brain processes involved in estimation of much more basic parameters such as distance itself. At what level this effect occurs could reveal the level at which the neural correlates of psychopathy are more dominant, something that could have clinical implications.

### Distance estimation bias

The perception of distance is instrumental in the recreation of three-dimensional information in the brain from the two-dimensional visual information produced by the retinae. It is largely determined by the processing and integration of both monocular and binocular cues of the input. The binocular cues are mostly informative for nearby objects where the images of the same object significantly vary between the two eyes. For objects located farther away, the binocular images are identical, and thus mostly the monocular cues are more informative for recreating depth information.

Recent research in humans has studied the brain correlates of distance perception when three-dimensional information is extracted mostly from either binocular or monocular cues (Berryhill & Olson, 2009). The main finding was that bilateral early visual areas V3A/B in the superior occipital gyrus have activations that are highly correlated with perceived distance. Most importantly, these areas appear to represent perceived distance even when distance estimation is task-irrelevant. These results provide strong evidence that distance information is automatically processed in early stages of visual processing. Deficits in distance estimation can be easily behaviorally quantified and measured. For this reason, this brain process is a good candidate for probing whether the neural correlates of psychopathy affect fundamental processes such as early visual feature extraction.

It has already been shown that some conditions involving irregular processing of emotions by the brain have an effect on distance estimation. Interpersonal distance estimation bias was found in non-clinical social anxiety in experiments of Givon-Benjio and Okon-Singer (2020). Individuals with social anxiety are characterized by avoidance of social interactions, manifested by a preference for greater interpersonal distance – specifically from strangers (Asnaani et al., 2014; Perry et al., 2013; Cohen et al., 2018; Clark & Wells, 1995). In order to find out whether the preference for distance is associated with estimating the physical interpersonal distance in a distorted manner, people with and without social anxiety were asked to participate in a computerized comfortable interpersonal distance task. Results demonstrated that socially anxious individuals estimate the interpersonal distance from strangers as shorter, so that the strangers seem to be in closer proximity. Biased distance estimation was also investigated with regard to acrophobia (Teachman et al., 2008). Participants were divided into high and low acrophobia groups based on their symptoms. In a visual matching task, participants viewed the vertical extent from a two-story, 26-foot high balcony with a target disk placed on the ground beneath the balcony. Participants estimated the height of the balcony by positioning an experimenter to be the same distance from them along the balcony as the participants were to the target on the ground. Although both groups overestimated vertical heights, the degree of overestimation was exaggerated in the high acrophobia group.

Kim and Son (2015) investigated how facial expression of emotion affects distance estimation itself. In this work the photographs employed showed actors with threatening expressions (anger, hate), neutral expressions (shame, surprise), or so-called safe expressions (pleasure, joy). Participants had to perform a visual matching task to report the perceived distance from each face. The study took place in a dark room where they were shown faces on a tablet computer simulated to be at various distances. To complete the task, participants first turned their heads 90° to the left to view the face on the tablet and then looked straight ahead to move a fluorescent panel to the perceived distance by manipulating a rope. Irrespective of emotions displayed, faces were perceived as closer than they actually were. Results showed that facial expression influenced distance estimation. Faces exhibiting threatening or safe expressions were judged significantly closer than those showing neutral expressions. Female participant’s judgements were more likely to be influenced by emotional face expression. The extent of underestimation by female participants was particularly pronounced for threatening expressions. The study also showed that there was a significant interaction between distance and facial emotion. In distance perception studies (Baird & Biersdorf, 1967; Johnston, 1991; Norman et al., 1996), distance underestimation generally increases with distance up to 75m (Hecht & Daum, 2009). In the research of Kim and Son (2015) a different pattern was found. For the safe and threatening expressions, the amount of underestimation grew with increases in distance, but for the two neutral expressions the amount of underestimation was greater at 2 m than at 1 m or 3 m.

### Aims of the present study

Psychopathy level has been shown to modulate the distance from another person at which someone feels uncomfortable according to the type of facial expression (Welsch et al., 2018). When asked for a comfortable preferred distance, people with low psychopathy index stopped farther away from an avatar with angry facial expression than when they approached an avatar with a happy face. In contrast, people with high psychopathy index stopped at a similar distance for both of these facial expressions. This was attributed to impaired processing of facial emotional content by people with high psychopathy index. At what level this effect occurs could reveal the level at which the neural correlates of psychopathy are more dominant, something that could have clinical implications. According to the dominant trend such an effect is attributed to higher processes, such as the processing of emotional content. Little attention however has been paid to possible low level processes that are affected by psychopathy, such as the extraction of basic physical parameters from sensory information. Here we aim to study distance estimation as one of these basic parameters. The information necessary for distance estimation by the brain is extracted from binocular (3D) or monocular (2D) cues, similar to the previously mentioned research of Berryhill and Olson (2009). In the current work we employ two-dimensional avatars at different distances and thus, we expect that the distance estimation is performed automatically in the same early visual processing areas V3A/B. This assumption has direct implications for forming the hypothesis as well as for interpreting the behavioural results of the current experiment. If distance estimation is affected by psychopathy level (measured behaviourally), then this indicates that the brain correlates of psychopathy affect very basic early stages of information processing in early visual cortex. This is not trivial as psychopathy is typically assumed to be related to brain processes higher in the cortical hierarchy, related more to the processing of emotional content and to emotion regulation.

In previous work it has been shown that in general the estimation of distance is affected by facial emotions (Kim & Son, 2015). In this work the distance to a face with high valence, positive or negative, was more underestimated than to a face with a neutral expression. This is evidence that facial emotional content has an effect on lower brain processes involved in the estimation of basic environment parameters such as distance. Exploiting this previous work, we decided to examine if this effect of facial expressions on distance estimation is modulated by psychopathy level. If the brain correlates of psychopathy do not reach early stage processing, then we would expect distance estimation to be affected by facial expression in the way demonstrated by Kim and Son (2015), irrespectively of the psychopathy level of the participants. Hence, for all psychopathy levels, distance to avatars with happy and angry face expressions would be underestimated because the distance is perceived as smaller than it actually is. However, if psychopathy affects early stage processing in the brain, we would expect results to show similar effects as previously found by Welsch et al. (2018). Hence, people with low levels of psychopathy would stop avatars at different distances, depending on their facial expression, while people with high level of psychopathy would have a similar distance estimate, independent of emotional facial expression. In such case, the closer avatar proximity for high psychopathy individuals would mean that in this context they overestimate distance (i.e. if they stop the avatar at 2.5 m when they were asked to stop it at a target distance of 3 m, they perceive 2.5 m of actual distance as 3 m). This finding would also have implications of clinical interest, as the pattern of distance estimation for different facial emotions could be used as a proxy for predicting the psychopathy level of an individual.

## Methods

### Participants

We recruited participants via the website and the database of the Max Planck Institute for Empirical Aesthetics in Frankfurt (Main) and via the email distribution list of the psychology student union at Johannes Guttenberg-University of Mainz. Participants filled in the questionnaires online. They were completed by 336 participants with an age range of 18 to 77 years (*M* = 30.54, *SD* = 11.67). Of these 251 were female, 82 male, one selected “other”, and 3 participants declined to answer. We invited all participants who filled in the questionnaire again for the online experiment. The sample for the main experiment consisted of 128 participants with an age range of 18 to 72 years (*M* = 30.61, *SD* = 12.41) with a majority of female participants (67 %). Based on their scores in the PPI-R, we split participants into three groups with balanced numbers per group: low, middle and high psychopathic individuals. Using three percentiles (divided at 33.3 % and 66.6 %), we labeled participants with a psychopathy score of 81 or lower as low, participants with a psychopathy score of 92 or higher as high, remaining participants as middle. These cut-off values created three groups with 42 (low), 43 (middle) and 43 (high) psychopathic individuals respectively. Assigning participants to the groups was not independent of participant gender, as indicated by a significant effect of interdependence, *χ*²(4) = 13.72, *p* < .008. Female participants were the majority in each group (low: 36 (85.7%), middle: 28 (65.1%), high: 22 (51.2%)) with significant overrepresentation in the low psychopathy group.

### Stimuli

#### Database

The emotional faces used for the experiment are from the “Karolinska Directed Emotional Faces” database (Lundqvist, Flykt, & Öhman, 1998). The actors who had been used for this database were between 20 and 30 years old. For the current study, we selected a set of 48 actors (24 male and 24 female), each with the face expressions happy, angry, and neutral.

### Animation and Online Deployment

The main question of the experiment regarded the estimation of distance from another individual. This dictated the need for a realistic representation of distance and for the presence of another individual. Therefore, we designed the experimental stimuli in three-dimensional space, rather than in two dimensions, so that the sense of depth enhanced the sense of distance estimation. We used two widely used animation software programs for the design of the stimuli in three-dimensional space, Blender and Unity. The online deployment was performed through a JATOS server.

#### Blender

We used Blender for the design of the animated character. We created a simple box-shaped avatar. The character was rigged with a basic human skeleton template of Blender. The head of the avatar was a cuboid. The face images from the KDEF-database were attached as textures to the front face of the head cuboid. The translational movement of the character was animated in blender. The avatar was animated to have a simple translation of 5.5 meters from its initial position. We decided to avoid any walking movements of the legs and arms in order to avoid the risk that the participants estimate distance by counting leg or foot movements. The translation of the avatar was performed with a speed of 1 m/s. We changed this initial speed later to two different speeds employed in the main experiment. In addition to the animated character and its simple translational movement, we used Blender to design a fixation mark, a panel for displaying the target distance to the participant, and a panel for providing feedback to the participant.

#### Unity

We designed the experiment in Unity, one of the most popular game design software programs, based on the C# programming language. The animation components built in Blender were imported into Unity where they were placed in the default 3D environment / background. We placed a default camera and light source placed at 5.5 meters away from the initial position of the animated character. The animated translation was also imported into Unity and was directed towards the camera. The camera view was used as the first-person view of the participants. We added a plane inside Unity, covering the upper part of the screen, which we used to give instructions to the participants. The sequence of stimuli with corresponding details was read from Unity from an ASCII file and placed inside the “Resources” folder of the project directory. There was a different ASCII file for every participant corresponding to a unique stimulus sequence. After we had designed and tested the entire experiment in Unity, we compiled it in a WebGL-compatible format so it could be run online on the participant’s preferred web browser. All participants received an email with the link to a web-server hosting the WebGL-version of the experiment. Once the participant had entered the link in the web-browser, the experiment was downloaded and run inside the browser. The participant’s responses were uploaded from her / his browser to the server hosting the experiment.

#### JATOS

JATOS (“Just Another Tool for Online Studies”) is an open-source server-side (back-end) tool, specifically developed to help researchers set up and run online experiments on their own servers. One main advantage is that it offers data security to the researchers as everything is run and stored on a server fully under their control. Another is that it comes with built-in participant management and recruitment tools. The Max Planck Institute for Empirical Aesthetics has installed a JATOS server so that its researchers can perform online experiments. The WebGL-version of the experiment was imported into the JATOS server, and then a common weblink was created which pointed to the experiment. Multiple participants could connect to the JATOS server and do the experiment in parallel. The JATOS API library is written in JavaScript and offers various functions, which can be used to upload data of the participant’s responses from the web browser running the experiment to the JATOS server from which it was launched. UNITY, in which scripting is exclusively in C#, offers a way to call such JavaScript functions by creating some intermediate JavaScript functions inside a .jslib-file in the “Plugins” subfolder of the assets folder of the experiment.

### Questionnaires

#### PPI-R

To measure psychopathy, we used a short version of the Psychopathic Personality Inventory-Revised (PPI-R) (original version: Lilienfeld, Widows, & Staff, 2005; German translation: Alpers, & Eisenbarth, 2008; short version: Eisenbarth, Lilienfeld, & Yarkoni, 2015). The short version of the PPI-R is a self-assessment questionnaire that consists of 40 statements, which have to be rated on a 4-point rating scale with the categories wrong, rather wrong, rather correct or correct. The scale has eight subscales, of which seven can be merged into two factors. The subscales social influence, stress immunity, and fearlessness can be assigned to a first factor that reflects Fearless Dominance. The second factor reflects Self-Centered Impulsivity and includes the subscales machiavellian egocentricity, rebellious nonconformity, blame externalization, and carefree nonplanfulness. There is also the subscale coldheartedness, which cannot be assigned to one of the two factors. The normalization of the questionnaire is based on a non-forensic student sample but can also be used for an older population (Alpers & Eisenbarth, 2008). The scores of all subscales were summed up to generate an overall psychopathy score, with higher scores indicating higher extent of psychopathy.

### Design and procedure of the main experiment

The main experiment consisted of two parts. Participants first had to rate face expressions, and subsequently they had to complete a distance estimation task after a training session. The duration of the whole experiment was around 30 minutes. As compensation, participants could choose between 10 Euro being transferred to their bank account and a lottery for a 60 Euro voucher for a bookstore. For participating in the experiment, the use of a computer or laptop instead of a tablet computer or smartphone was mandatory.

#### Rating face expressions

Participants were asked to rate a randomly selected set of 24 faces from the actual experiment (8 happy, 8 neutral and 8 angry face expressions). The set was different for each participant. The ratio of male and female actors in the rating set was equal. The rating scale consisted of 9 points, ranging from “negative” to “positive”. The middle rating point was labeled as “neutral” (see Figure 1).

**Figure 1.**
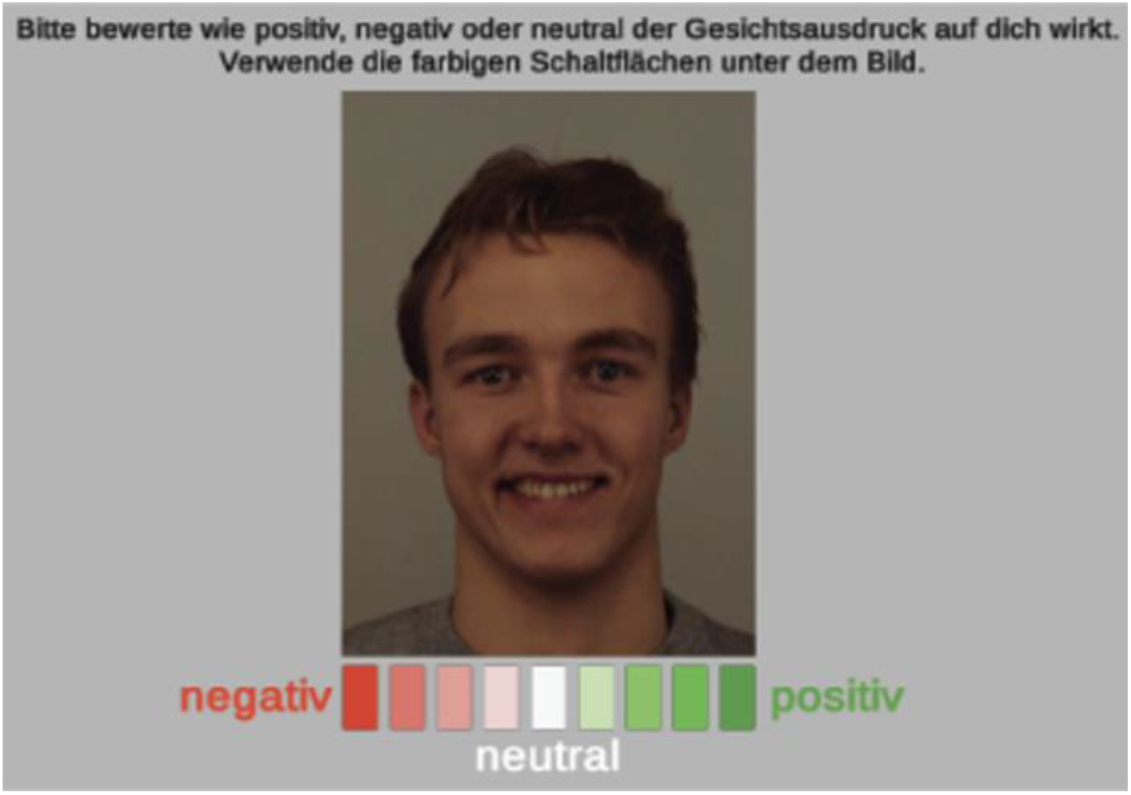
Rating of face expressions. Participants had to select a mark on a scale ranging from negative (red) to positive (green).

#### Distance estimation

In the main task, we asked participants to estimate a target distance between an avatar and themselves. In the beginning of each trial, the target distance was shown in the middle of the screen as a decimal number in meters (Figure 2). The starting distance of the avatar was always 5.5 m. The height of the avatar was 1.73 m. After a button press by the participant, the target distance display disappeared and the avatar appeared at the starting distance with a fixation marker. After 0.5 seconds the fixation marker disappeared and the avatar started “walking” towards the participant. The walking speed of the avatar was either slow (1.25 m/s) or fast (1.75 m/s) in 50 % of the trials, respectively. The participant’s task was to stop the avatar in the previously displayed target distance as accurately as possible. In the instruction, participants were frequently reminded to focus on the avatar’s face at any time.

**Figure 2.**
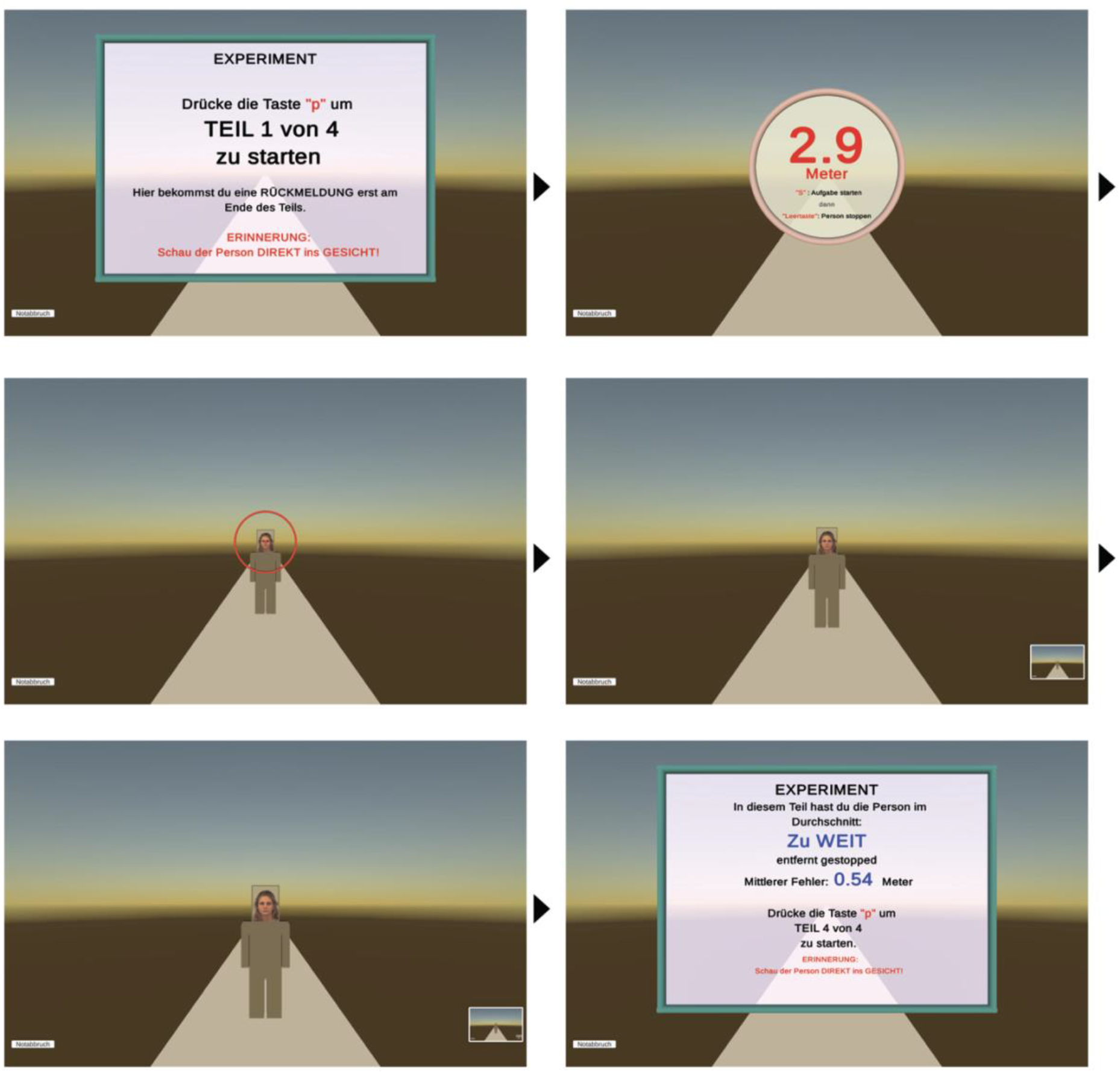
Distance estimation task. At the beginning of each trial the participant was presented with a target distance where the avatar should be stopped. The task starts with a participant button press. The avatar was presented in its initial position and a fixation marker (red ring with small sphere in center) was used to focus the participant’s attention on the face of the avatar. The avatar started moving towards the participant, who had to stop it by pressing the space bar at the previously displayed target distance. With the participant’s button press, the avatar stopped instantly. The participant had to press a button to proceed to the next trial. At the end of each of the 4 blocks, the participants received feedback about their average error in distance estimation. Please note that feedback was not given after individual trials.

As target distance there were six different distances ranging from 0.5 m to 3.6 m. In order to avoid the situation where the subjects are always presented with the same set of six distances and stopped paying attention to the displayed distance, we introduced variability by having two different sets of distances. Distance set A ([0.5, 1.1, 1.7, 2.3, 2.9, 3.5] meters) and distance set B ([0.6, 1.2, 1.8, 2.4, 3.0, 3.6] meters) varied within 0.1 m for each distance. One distance set was always used for the slow speed trials and the other for the fast speed trials. The exact type of distance set allocated to each speed was randomized across subjects. For half the subjects the allocation was distance set A – slow speed, and distance set B – fast speed. For the other half of the subjects the allocation was vice versa.

We divided the whole experiment into four blocks, each block consisted of 36 trials. This resulted in 144 trials for every participant. We balanced the number of emotional face expressions per block. After each block, participants received a summary feedback about their error as a decimal number. Therefore, the average error over all trials of this block was calculated. Participants did not receive a feedback after each trial. If the avatar had been stopped within a range of +/- 0.5 m on average in all trials of the block, the feedback was labeled as “perfect” (“genau”). If the avatar had been stopped closer or farther than 0.5 m to the target distance, participants received the feedback “too close” (“zu nah”) or “too far” (“zu weit”), respectively. For analysis, a positive error value referred to underestimation (the avatar was stopped too early / far) and a negative error value referred to overestimation (the avatar was stopped to late / close) as described in figure 3.

**Figure 3.**
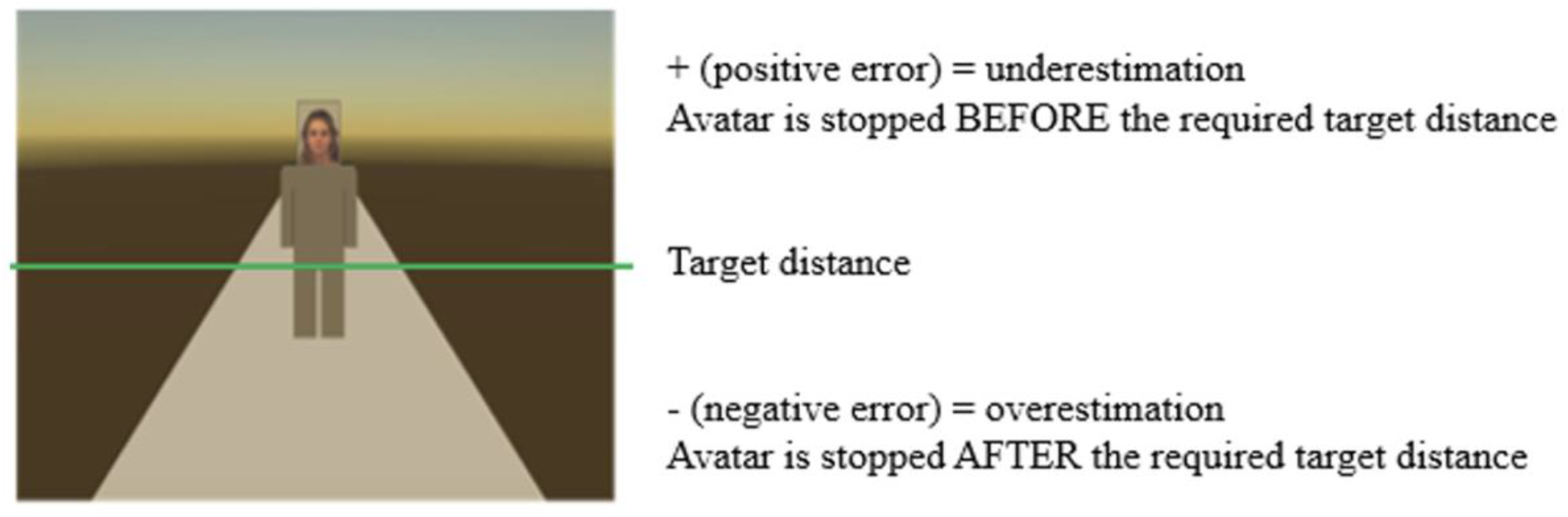
Meaning of error values. For analysis, positive errors reflect underestimation, indicating that the avatar appeared closer than it actually was and was therefore stopped too early. Negative errors reflect overestimation, indicating that the avatar appeared farther away than it actually was and was therefore stopped too late.

Overall, three within-subject variables were controlled in the experiment: emotional expression, speed, and distance. Thus, the experiment utilized a 3 (emotion: neutral, angry, happy) x 2 (speed: slow, fast) x 6 (distances from 0.5 m to 3.6 m) mixed-design. This resulted in 36 different combinations of emotion-speed-distance. For each combination, there were 4 trials, each one with a different actor image. Consequently, there were in total 144 trials per participant.

#### Training

Participants were prepared with a training prior to the main experiment to make them feel comfortable with the buttons they had to press. An avatar with a neutral face expression who was not part of the main experiment was used. Like in the main experiment, participants had to estimate a previously shown target distance between an avatar and themselves. The sequence of events in each trial of the training session is identical with the sequence of events in trials of the main experiment. The target distance was displayed at the beginning of the trials. When the participant started the trial, the target distance display disappeared and the avatar appeared at the starting distance of 5.5 m. After 0.5 seconds, the avatar started “walking” towards the participant. Unlike in the main experiment, the walking speed of the avatar in the training was always slow (1.25 m/s). The participant’s task was to stop the avatar in the previously shown target distance as accurately as possible. The training session consisted of 40 trials and could be repeated by the participant if desired. Whereas in the main experiment, participants received feedback only after each block, in the training session feedback was given after each trial. The target distance for the training was 0.5 m, 1 m, 2 m, 3 m or 4 m. Each of these distances was used in 8 trials. The sequence of presentation of the training trials was presented in the same order for all participants.

### Data Analysis

When the ANOVA sphericity assumption was violated (Mauchly’s test, *p* < .05), we used the Greenhouse–Geisser correction (Winer, 1971). The fractional degrees of freedom indicate this correction. All analyses were performed using SPSS for Windows (version 25). Partial *η*² is reported as an estimate of effect size in the rmANOVA.

## Results

### Ratings

Participants had to rate a randomly assigned set of 24 faces, with balanced number of emotional face expressions. The range of possible ratings was -4 to +4. For each of the emotional face expressions (neutral, angry, happy), mean ratings were averaged for each subject. For testing if psychopathy level had an effect on the rating of the emotional faces, mean ratings were entered as dependent variable into an analysis of variance (ANOVA) for analysis with repeated measure on the factor emotional face expression. Psychopathy group was entered as between-subjects factor. Prior to the ANOVA, Shapiro-Wilk tests were performed for mean ratings of all three conditions to test whether assumption of normality was valid. Results were significant in all cases, indicating that the data was not normally distributed, *p* < .001. However, because of similar sample sizes, violation of the normal distribution assumption was accepted. There was homogeneity of the error variances, as assessed by Levene’s test (*p* > .05).

The ANOVA confirmed a main effect of emotion, Greenhouse–Geisser *F*(1.5, 187.0) = 2020.87, *p* < 0.001, *η*² = .94, but not for psychopathy group, *F*(2, 125) = 0.82, *p* = .442. Neutral face expressions were on average rated with -0.46 (95 % - CI [-0.54, -0.37]), angry face expressions had an average rating of -2.89 (95 % - CI [-2.99, -2.79]) and happy face expressions received an average rating of 2.45 (95 % - CI [2.28, 2.61]). The results show that all participants, independently of the assigned psychopathy group, were able to recognize the emotional content of the face expression correctly. We did not find any other main effect or interaction effect.

### Distance estimation

For every trial, the distance estimation error was calculated by subtracting the target distance from the distance at which the participant actually stopped the avatar. Outliers were determined in three steps. First, the optional comments of all participants at the end of the experiment were reviewed and possible problems with the correct presentation of the stimuli were identified. Four participants reported complications during the distance estimation task and were therefore excluded from further analysis. Subsequently, we searched for participants who had estimation errors of more than 1 m in more than 50 % of all trials. This applied to one participant who therefore was also excluded. Last, all trials with error values larger than 1 m were discarded. For 13 participants this resulted in some of the emotion-speed-distance combinations having no trials. For this reason, these 13 subjects were excluded. The remaining sample included 110 participants (18 - 70 years, *M* = 29.6, *SD* = 11.5) of which the majority was female (67.3 %).

These participants were grouped into low, middle and high psychopathic groups by using the 1/3 and 2/3 percentile values of the overall score of the PPI-R questionnaire. 39 participants with a score of 81 or lower were assigned to the low psychopathy group, 35 participants with higher scores until 89 to the middle psychopathy group, and 36 participants with scores higher than 90 to the high psychopathy group.

Assigning participants to the groups was again not independent from participant gender, indicated by a significant effect of interdependence, *χ*²(2) = 17.4, *p* < .001. Most notable is that female participants are significantly underrepresented in the high psychopathy group (distribution of female participants per psychopathy group: low: 35 (89.7%), middle: 23 (65.7%), high: 16 (44.4%)).

To reveal the imbalance of male and female participants regarding distribution of psychopathy score, we sorted participants according to their score. Figure 4 shows a higher number of male participants for high psychopathy scores and a higher number of female participants for low psychopathy scores.

**Figure 4.**
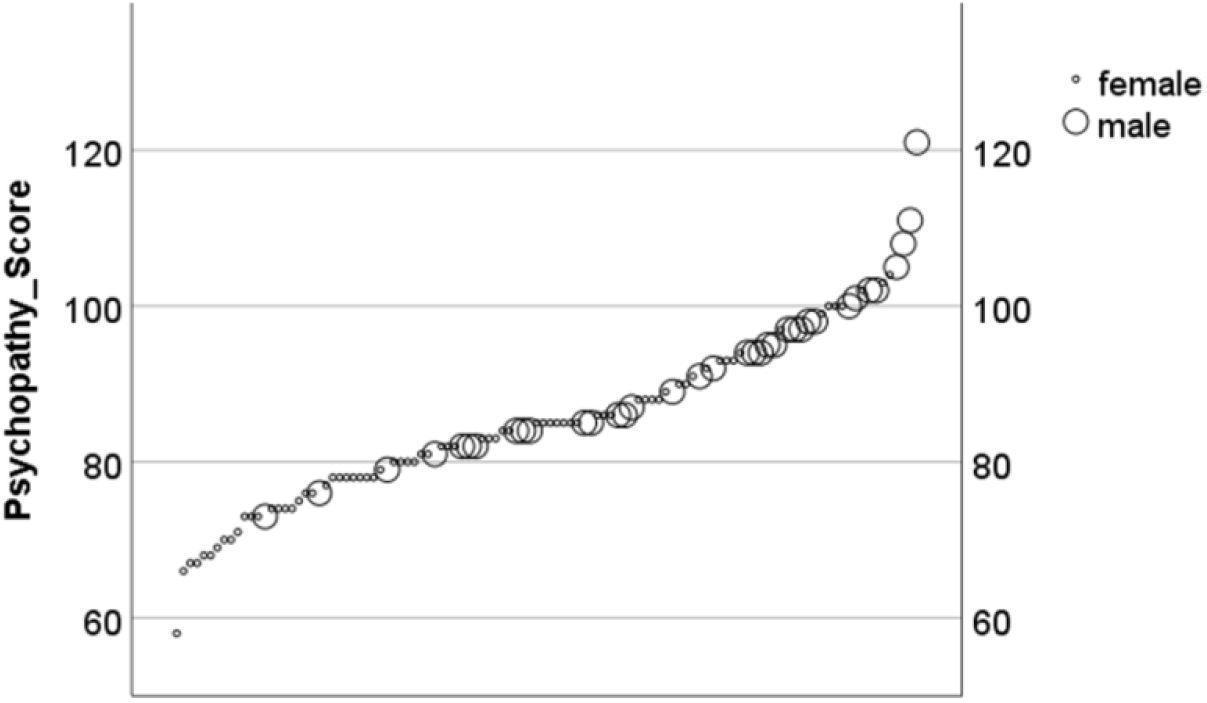
Sorted scatterplot of psychopathy score and participant gender.

We entered distance estimation error as dependent variable into a mixed-design analysis of variance (ANOVA) for analysis with emotional face expression, speed, and distance as independent variables. Psychopathy group was entered as between-subjects factor. There was homogeneity of the error variances, as assessed by Levene’s test (*p* > .05). Prior to the ANOVA, we also performed Shapiro-Wilk tests on dependent variables to test whether assumption of normality was valid. Results were significant, indicating that the data is not normally distributed, *p* < .05. However, repeated measure ANOVA is relatively robust against violations of the normal distribution assumption (Pagano, 2010; Salkind, 2010; Wilcox, 2012), therefore no data transformation was performed prior to the analysis.

The ANOVA confirmed a main effect of psychopathy group *F*(2, 107) = 3.74, *p* < 0.05, *η*² = .07. Participants with low and middle psychopathy scores showed similar estimation errors of 14.7 cm (95 % - CI [9.7, 19.7] cm) and 16.1 cm (95 % - CI [10.8, 21.4] cm), respectively. However, high psychopathic individuals showed significantly lower errors with 6.6 cm (95 % - CI [1.4, 11.9] cm) on average (see Figure 5). Bonferroni-adjusted post-hoc analysis revealed a significant difference of distance estimation errors (*p* < .05) between the middle psychopathy group and the high psychopathy group (9.5 cm, 95 % - CI [0.3, 18.6] cm).

**Figure 5.**
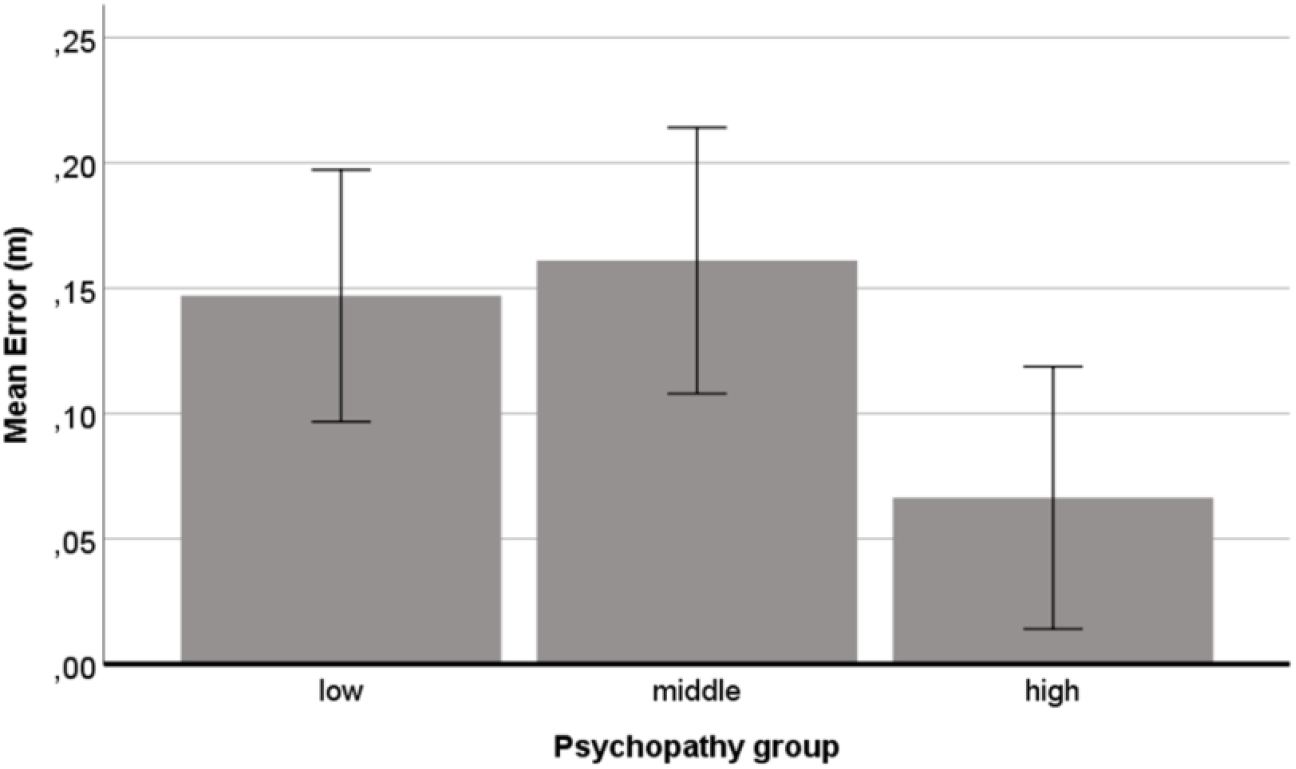
Mean distance estimation error in meters per psychopathy group. Positive errors reflect underestimation. Error bars represent the 95% confidence interval. *p < .05.

Statistically significant differences of the distance estimation error were also found for the factor Distance, Greenhouse–Geisser *F*(2.4, 260) = 24.80, *p* < .001, *η*² = .19, and the factor Speed *F*(1, 107) = 166.25, *p* < .001, *η*² = .61. Figure 6 shows mean errors of distance estimation for the slow and fast speed condition. Table 1 shows pairwise comparisons of mean distances in meter to identify significant differences.

**Figure 6.**
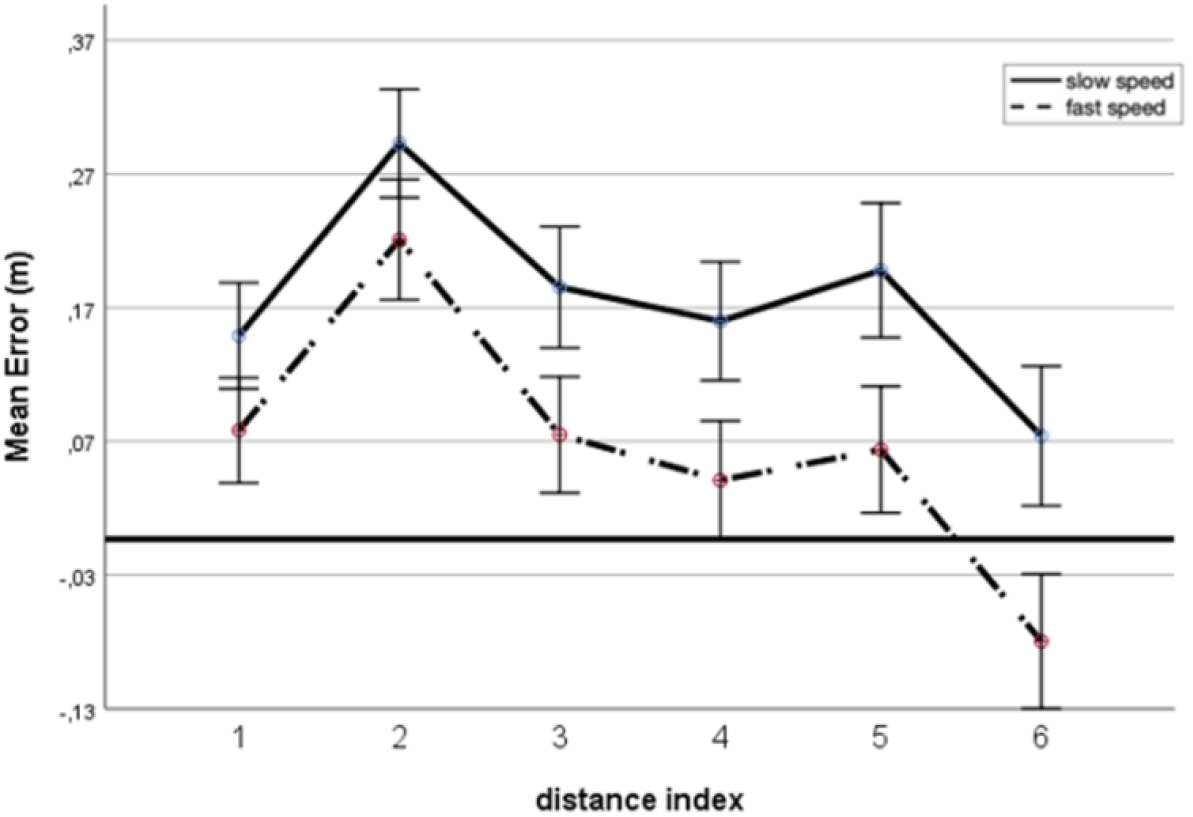
Mean distance estimation error in meter per distance. Positive errors reflect underestimation. Distance indices: 1: [0.5 or 0.6] m, 2: [1.1 or 1.2] m, 3: [1.7 or 1.8] m, 4: [2.3 or 2.4] m, 5: [2.9 or 3.0] m, 6: [3.5 or 3.6] m. Error bars represent the 95% confidence interval.

**Table 1.** Pairwise comparisons of mean distances in [m]

| Slow speed |  |  |  |  | Fast speed |  |  |  |  |  |  |
| --- | --- | --- | --- | --- | --- | --- | --- | --- | --- | --- | --- |
|  | 1 | 2 | 3 | 4 | 5 |  | 1 | 2 | 3 | 4 | 5 |
| 1 | - |  |  |  |  | 1 | - |  |  |  |  |
| 2 | -.432* | - |  |  |  | 2 | -.426* | - |  |  |  |
| 3 | n.s. | .325* | - |  |  | 3 | n.s. | .441* | - |  |  |
| 4 | n.s. | .402* | n.s. | - |  | 4 | n.s. | .544* | n.s. | - |  |
| 5 | n.s. | .287* | n.s. | n.s. | - | 5 | n.s. | .474* | n.s. | n.s. | - |
| 6 | n.s. | .656* | .331* | .254* | .369* | 6 | .473* | .900* | .459* | .356* | .426* |

Note. Mean differences of mean distances in [m] across both speed conditions.
Distance indices: 1: [0.5 or 0.6] m, 2: [1.1 or 1.2] m, 3: [1.7 or 1.8] m, 4: [2.3 or 2.4] m, 5: [2.9 or 3.0] m, 6: [3.5 or 3.6] m. $N = 110$ . Bonferroni-corrected, $*p < .05$

Figure 7 shows the estimation error as a percentage of the correct distance. There was underestimation for all distances, which means that the avatars had been perceived closer than they actually were or in other word avatars had been stopped too early. In the slow speed condition, the extent of underestimation was highest for the closest distance (0.5m or 0.6m) and decreased with distance with the farthest distance leading to the lowest extent of underestimation. For the fast speed condition a similar pattern was found but with an overall lower extent of underestimation. In the slow speed condition, distance estimation errors were 11cm larger (*p* < .001) than in the fast speed condition (11.0 cm, 95% - CI [9.3, 12.7] cm), as shown in figure 8. The approaching avatars were stopped significantly earlier in the slow speed condition. In both speed conditions avatars were stopped too early / too far away.

**Figure 7.**
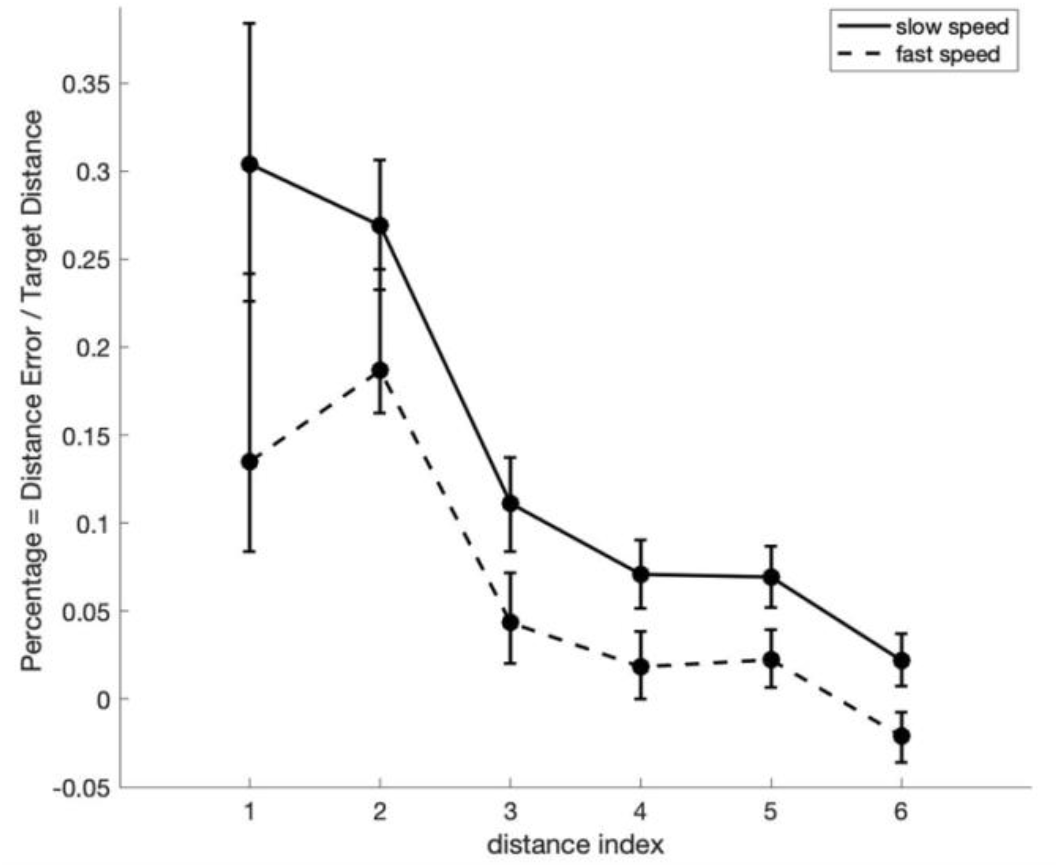
Ratio of distance errors over target distance per distance. Positive errors reflect underestimation. Distance indices: 1: [0.5 or 0.6] m, 2: [1.1 or 1.2] m, 3: [1.7 or 1.8] m, 4: [2.3 or 2.4] m, 5: [2.9 or 3.0] m, 6: [3.5 or 3.6] m. Error bars represent the 95% confidence interval.

**Figure 8.**
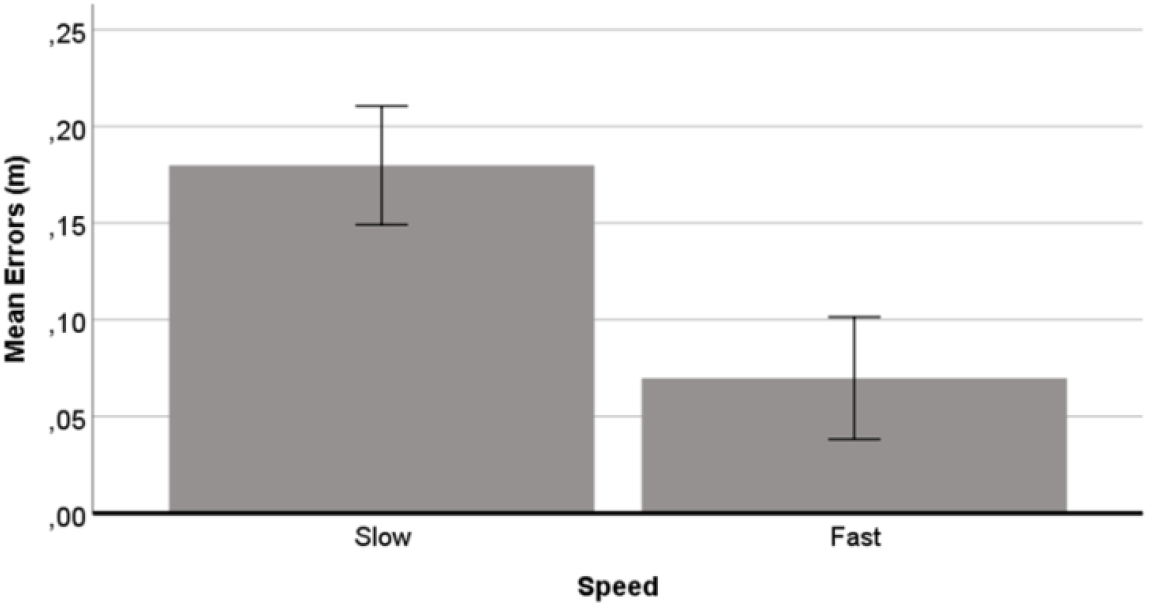
Mean distance estimation error in meter per Speed condition. Positive errors reflect underestimation. Error bars represent the 95% confidence interval.

The factor Emotion did not reach statistical significance, *F*(2, 214) = 0.17, *p* = .846, indicating that emotional face expression had no effect on distance estimation, regardless of psychopathy group.

To test for the impact of the subscales of the overall PPI-R score, we conducted a rmANOVA with subscale scores as between-subjects variable. The scores of the subscales Self-Centered Impulsivity and Fearless Dominance were each divided into three groups. This resulted in approximately equal numbers of participants in each group (Self-Centered Impulsivity: 39 low (score 0 to 37), 34 middle (score 38 to 42), 37 high (score 43 or more); Fearless Dominance: 39 low (score 0 to 34), 37 middle (score 35 to 39), 34 high (score 40 or more)). Since the facet Coldheartedness did not load on the other subscales and therefore had a lower range of possible scores, participants were grouped into only two groups with low and high extent of Coldheartedness (47 low (score 0 to 9), 63 high (score 10 or more)). Neither Coldheartedness, *F*(1, 108) = 0.01, *p* = .935, nor Fearless Dominance reached statistical significance, *F*(2, 107) = 0.37, *p* = .691. However, for the subscale Self-Centered Impulsivity a significant difference was found, *F*(2, 107) = 4.01, *p* = .021, *η*² = .07. As there was violation of homogeneity of the error variances, as assessed by Levene’s test, Games-Howell post-hoc analysis was performed. This revealed a significant difference (*p* < .05) between participants of the low and high Self-Centered Impulsivity group in their distance estimation errors (10.26 cm, 95% - CI [0.98, 19.5] cm). Figure 9 shows that participants with high scores on Self-Centered Impulsivity let the avatar approach significantly closer than participants with low scores.

**Figure 9.**
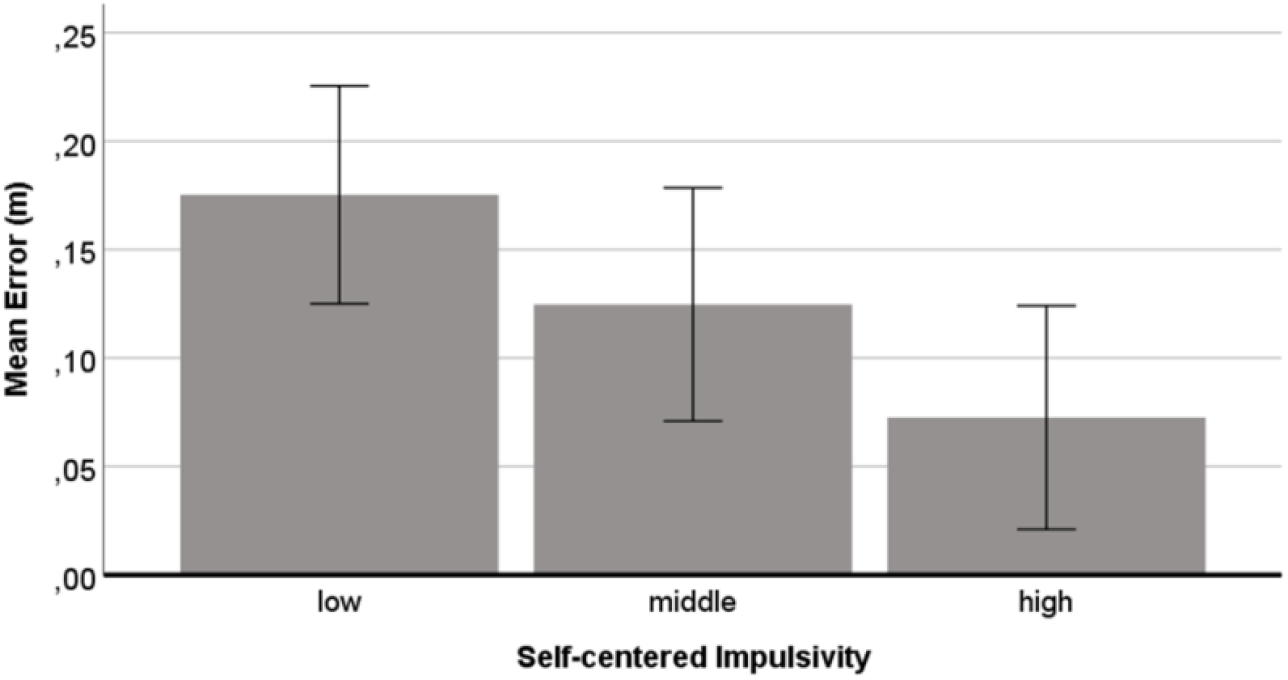
Mean distance estimation error in meters for low middle and high level of Self-Centered Impulsivity. Positive errors reflect underestimation. Error bars represent the 95% confidence interval.

Additionally, we also conducted a rmANOVA separately with all five dimensions of the BFI-10 as between-subject variable. None of the dimensions showed significant effects on distance estimation errors (*p* > .05).

## Discussion

The main goal of this thesis was to investigate the hypothesis that psychopathy has an effect on elementary processes in the brain such as the estimation of distance. Distance estimation can be considered an early-stage process and has been shown by recent research to be affected by emotional context (Kim & Son, 2015). For the present study, we first invited volunteers to participate in an online questionnaire. According to the questionnaire results, we divided participants into low, middle, and high psychopathic individuals. Subsequently a representative subset of participants performed an online virtual distance estimation task. In this task participants had to stop an approaching avatar with different facial expressions at various target distances. We found that all participants underestimated the distance of the avatar, irrespectively of the avatar’s emotional face expression. The extent of underestimation was significantly smaller for participants with high levels of psychopathy as compared to participants with low or moderate levels of psychopathy. This is a key finding, demonstrating that psychopathy has an effect on fundamental feature extraction in early visual processing.

According to the initial hypothesis we expected, based on the previous work of Welsch et al. (2018) and Kim and Son (2015), that people with a low level of psychopathy would stop angry avatars farther away than happy avatars, whereas people with a high level of psychopathy should stop both at similar distance. Another expectation based on previous work by Kim and Son (2015), was that in general, participants would stop high valence avatars (happy and angry face expression) farther than avatars with neutral expressions. Our results did not confirm any of the two above expectations, as we did not find any effects of emotion on distance estimation.

### Emotion

Regarding an effect of emotional face expression on distance estimation accuracy, we failed to replicate the findings of Kim and Son (2015). Because in our results the manipulation of the emotion of face expression had no effect on distance estimation, we could not find the expected interaction effect between emotion and psychopathy. Independent of their psychopathy score, participants showed no differences in distance estimation as a function of the facial expression of the avatar. Possibly one of the main reasons that the replication was not successful were different experimental procedures. Looking closer into the experimental setup of Kim and Son (2015), the task was very different in many aspects from the task in the present study. Participants in the setup of Kim and Son (2015) were less under time pressure when making their distance judgements. In addition, their experiment took place in a dark room without context information and an experimenter was always present. However, in the present study, the avatar started to approach the participant immediately at the beginning of each trial, giving participants less time to think about their judgment and no possibility for corrections after their response. Additionally, distance estimation on a computer within a virtual reality makes the task more difficult. Although participants were repeatedly instructed to focus on the approaching avatar’s face (to be confronted with the emotional expression), we cannot be sure how conscientiously the participants have adhered to this instruction.

### Distance

On average, interpersonal distance was underestimated in all conditions and independently of psychopathy level. Avatars were stopped too far, indicating that participants perceived them to be closer than they actually were. Also Kim and Son (2015) found this effect in their study. Furthermore, our data showed that estimation accuracy increased with distance in the tested range of 1.1 m to 3.7 m. This result is contrary to what Kim and Son (2015) found in their work but consistent with previous literature on distance perception (e.g. Baird & Biersdorf, 1967). We attribute this difference to the completely different experimental conditions employed by Kim and Son (2015), where face images where placed standalone on panels, which participants had to move manually by a string mechanism. The experimental environment of the current work was a virtual environment, presented on the screen of the computer with a simple landscape as a background and in which the face images were actually attached on an avatar. We assume that participants’ distance estimation accuracy is highly affected by the presence of cues in the environment. The study of Kim and Son (2015) does not provide any environmental cues for an enhanced accurate distance estimation as the study took place in a dark room. We believe that this is an important difference that can explain the different patterns of accuracy variation across distance to the present study.

### Psychopathy

We found a significant effect of psychopathy. High psychopathy individuals underestimated distance significantly less than did participants of the middle or low psychopathy groups. All psychopathy groups underestimated the distance, meaning that the average distance error was positive for all, with the high psychopathy group having the smallest mean error. One could argue that psychopaths are just more accurate in a distance estimation task. However, the fact that all psychopathy groups had a positive mean distance error, i.e. all underestimated distance, suggests that the experimental virtual environment had a bias, leading the participants in general to underestimate distance. A positive bias would mean that distance in the virtual world design of the experiment on the computer appears generally larger which would result in overall underestimation. The extent of the underestimation by the bias is unknown for the present study since there was no control. Without knowing the exact bias of the method, it is not possible to draw conclusions whether low or high psychopaths have actually better estimate of the actual distance. The initial aim of the study was to compare levels of low and high psychopathy and discuss the differences. To that extent, the main inference is that the high level psychopathy group underestimates distance to approaching avatars less (lets the avatar approach closer) as compared to the other psychopathy levels. But a suggestion for implementing a possible control in the study design will be discussed in the paragraph “Future research”.

Regarding comfortable interpersonal distance, Welsch et al. (2018) could not find this main effect when participants had to choose their comfortable interpersonal space. This might be explained by the fact that the participants had to assess comfort, a process which might include complex high level processes, as compared to the simple distance estimation task at hand in the current study. However, Vieira and Marsh (2014) showed that Coldheartedness predicted preferred interpersonal distance, with more coldhearted participants preferring shorter distances. Their finding is confirmed by our results. In the present study we also tested for the subscales of the PPI-R including Coldheartedness. We only found a significant difference for the subscale Self-Centered Impulsivity, but not for Coldheartedness or Fearless Dominance. Participants who scored high on Self-Centered Impulsivity had the smallest distance estimation error (let the avatar approach closer). Hence, our study was able to show that components of psychopathy certainly contribute to altered distance estimation processing.

A closer look into the interpretation of high scores in the subscales that are merged into the factor Self-Centered Impulsivity might explain this effect. It comprises four lower-order subscales that assess a narcissistic and reckless tendency to exploit and blame others (Benning et al., 2003). In the literature, Fearless Dominance is related to a lack of emotional responsivity but accurate perception of those emotions in others. Conversely, Self-Centered Impulsivity is related to difficulties in both emotional perception and control of negative emotional responses, such as anxiety, irritation, and aggressiveness (Del Gaizo & Falkenbach, 2008). Book (2005) suggests that people scoring high on Fearless Dominance are better at identifying emotional facial cues, specifically fear and anger, in order to be successfully deceitful or know when to change strategies. However, people with higher scores on Self-Centered Impulsivity who tend to be reactively aggressive and demonstrate hostile attribution biases, demonstrate more errors in emotional perception. Taking these findings together, one could discuss a possible explanation for the effect in the present study. Although in our data we could not find a significant interaction effect of emotion and Self-Centered Impulsivity, participants with higher Self-Centered Impulsivity level might be less distracted by the emotional face expressions in general. This could result in their tendency to have more accurate distance estimation independently of face expression. Also Welsch et al. (2018) found that participants who scored higher on Self-Centered Impulsivity regulated distance less according to facial expression. They argue that this could reflect the tendency of those participants to not integrate peripheral information of social cues into the own behavior when engaging in goal directed behavior as proposed by the response modulation hypothesis of psychopathy (Smith & Lilienfeld, 2015). Our data support this argument.

### Speed

We could observe a strong effect of the speed of the approaching avatar on distance estimation accuracy. In the slow condition participants tended to make larger errors. Hence, the avatars were stopped too early. In the fast condition, avatars were still stopped too early but to a smaller extent. One simple explanation for this observation could be that the slow speed makes participant impatient and they could not wait the appropriate time for stopping the avatar in the correct position.

We can also think of another possible explanation that involves Representational Momentum (RM). RM is a relatively small error in our visual perception of moving objects. It has been attributed to forward extrapolation of an object’s motion so that people think that an object is a bit further along its trajectory as time goes forward. A higher speed of an object leads to a larger error due to RM (Choi & Scholl, 2006). In our case this would mean that although in both the fast and slow speed conditions the extent of underestimation might be the same, due to the higher RM error in the fast condition the participants stopped the avatar slightly further along its path. This then could be manifested as a smaller underestimation of distance such as what our results show.

Another potential explanation could be formulated however, if it is assumed that the experimental procedures have in overall a positive bias in distance estimation. As mentioned earlier there was an average positive distance estimation error for all psychopathy groups and for all conditions (speed-emotion). This overarching positive tendency of distance estimation error hints that the experimental procedures (a 3D avatar displayed on a 2D monitor) has potentially a bias, leading participants in general to stop avatars at longer distances than the target. If we hypothesize that this is indeed the case then it could also be hypothesized that this bias is close to the average distance estimation error of the slow condition. And this is because people in the slow condition have enough time to form their prediction and stop the avatar correctly. In this case, after removal of such a bias, the average error for the fast condition would turn negative. This would mean that in this case the participants stopped the avatar too late, and it is only due to the bias that it appears as if there was a lower underestimation error for the fast condition. This explanation would fit well the simplistic expectation that a fast-moving avatar would “slip” closer to the participant as compared to a slow moving one, only due to the longer distance travelled for the same reaction time.

### Ratings

Prior to the main task of distance estimation, participants were asked to rate the valence of the avatar’s emotional face expressions. Literature provides evidence for impaired emotion recognition in psychopathic individuals, but only in either time-restricted recognition tasks with high psychopathic inmates (Book, Quinsey, & Langford, 2007; Habel et al., 2002) or only for specific emotions like fear or disgust (e. g. Iria & Barbosa, 2009; Hansen et al., 2008) that were not relevant for the present study. We found no significant differences between the different levels of psychopathy on rating the face expressions. Independently of the extent of psychopathy, participants recognized the neutral, happy and angry face expressions correctly. Thus, the data of the rating part in our experiment confirm that participants with higher psychopathy levels had no difficulties recognizing the emotional content of the avatar’s faces employed in the subsequent task.

### Distance compression

An important parameter that needs to be taken into account in the assessment of the results is that there is an apparent compression of distance error, which grows with distance. For example, the apparent avatar’s size to an observer is much more similar when the avatar moves from 3.5 m to 3 m as compared to the case when the avatar moves from 1 m to 0.5 m, in which case the apparent avatar’s height changes much more.

This is explained more intuitively in Figure 10 (a). In this figure one can see the observer’s field of view (FOV) represented by two diverging blue lines. For this demonstration, the FOV has been selected as 150 radians, a typical angle for the vertical human FOV. The vertical blue arrow lines represent the height of the FOV as different distances from the observer (the farther away from the observer the larger the height of the FOV). The green arrow represents the actual height of the avatar. When the avatar is positioned very near the observer, it occupies a large proportion of the FOV. As the avatar moves away from the observer, it progressively occupies less and less of the FOV. Here the proportion of the FOV occupied by the avatar at a given distance from an observer has been quantified for illustration purposes as the ratio of the height of the avatar divided by the height of the FOV at this distance. This ratio is plotted in figure 10 (b) for the 6 distances used in the experiment (0.5 m to 3.5 m). It is obvious that for far distances, above 2 m, the avatar appears to occupy very similar proportions of the FOV. For an observer it should be challenging to accurately distinguish between them, based on the apparent size of the avatar. For near distances, smaller than 2 m, the ratio of the FOV occupied by the avatar increases exponentially, the closer the avatar is to the observer. It should be easier for the observer to distinguish between different near distances of the avatar, based on its apparent size. According to the above expectations, participants should have higher distance estimation errors for the far distances, as they are more difficult to differentiate, and smaller errors for the short distances, as they can be distinguished more easily. Our results do not seem to support this hypothesis. The plots of the average estimated distance error per distance in Figure 6 do not show such a pattern. In contrast these plots show that the estimated distance error is very similar across most distances with a large positive deflection for distance index 2 (1.1 m and 1.2 m), where the error is largest. We find a big negative deflection for the farthest distance index 6 (3.5 m and 3.6 m), where the error is small. If the estimation error was strongly modulated by the compression of the apparent distance, as depicted in Figure 10 (b) then the estimated distance error should be highest at the farthest distances and lower at the near distances. One alternative hypothesis could be, as already discussed, that the average estimated distance error has an inherent methodological positive bias. If in such a case the bias would be around the average error of the near distances (distance index 1 and 2 in figure 6), then indeed the largest error would be for the farthest distances (distance index 6 in figure 6). However, this would not solve the fact that the shape of the estimated error with distance is very different from the shape of the distance compression with distance. Finally, as we have already stated, the current experimental design does not allow investigating whether there is a bias or not. So the only concrete conclusion that can be drawn from the current work is that the estimated distance error does not seem to be modulated by the compression of the apparent distance.

**Figure 10.**
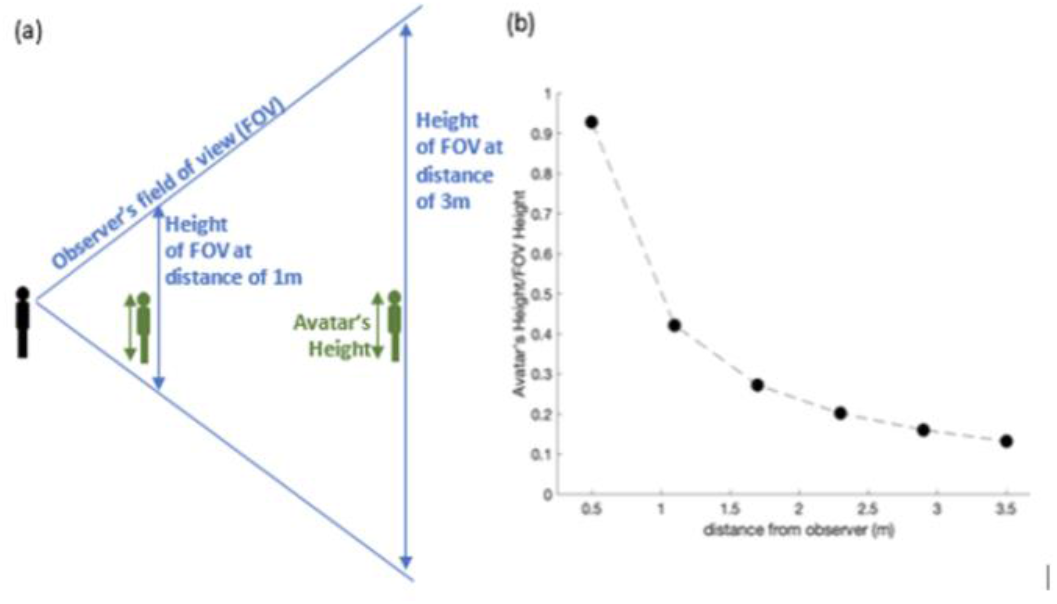
Distance compression. (a) Demonstration of the observer’s field of view (FOV) at an angle of 150°. The vertical blue arrow lines represent the height of the FOV as different distances from the observer (the farther away from the observer the larger the height of the FOV). The green arrow represents the actual height of the avatar. (b) Ratio of avatar’s height / FOV height vs. distance from observer in meters.

## Conclusion

Our work makes a significant new contribution to the study of psychopathy. It demonstrates through a simple distance estimation experiment that individuals with high psychopathy level underestimate distance of an approaching avatar with a human face less, irrespective of the emotional content of the expression, as compared to individuals with low or moderate psychopathy level. The significance of this finding stems from the fact that distance estimation is considered an automatic process performed in early stages of visual processing. The fact that psychopathy appears to affect such basic feature extraction in sensory cortex demonstrates that the neural correlates of psychopathy span the entire functional hierarchy of the brain. In terms of the effect of emotional face expression on distance estimation, we did not find any significant differences between individuals with low and high extent of psychopathy. This could mean that the effect of the valence of the facial expression does not reach as low as early visual processing, when it is not relevant to the task. These findings hint that it is the mere presence of another person, irrespective of the valence of emotional expression that is processed differently in the brains of low and high psychopaths.

## Notes

### Competing Interest Statement

The authors have declared no competing interest.

